# Tethered exosomes containing MT1-MMP contribute to extracellular matrix degradation during breast cancer progression

**DOI:** 10.1101/2024.10.29.620634

**Authors:** Roberta Palmulli, Hannah K. Jackson, James R. Edgar

**Affiliations:** Department of Pathology, University of Cambridge, Cambridge. CB2 1QP. UK; Exosis, Inc. Palm Beach, Florida. 33480. USA

**Author notes:** Corresponding author, JRE.

## Abstract

For cancer cells to escape from the primary tumor and metastasize, they must degrade and navigate through the extracellular matrix (ECM). The transmembrane protease MT1-MMP plays a key role in localized matrix degradation, and its overexpression promotes cancer invasion. In this study, we demonstrate that MT1- MMP is trafficked to the intraluminal vesicles of multivesicular endosomes, and subsequently released from cells on exosomes, a subtype of extracellular vesicle that can be retained to the surface of the originating cell by the anti-viral restriction factor, tetherin. While tetherin overexpression is linked to increased cell migration and invasion in various cancers, its role in these processes remains unclear. Our findings reveal that expression of tetherin by breast cancer cells promotes the retention of MT1-MMP-positive exosomes at their cell surface, while tetherin loss enhances exosome escape and impairs ECM degradation. Thus, tethered exosomes promote the retention of MT1-MMP at the surface of cells, aiding the degradation of the ECM and promoting cancer cell invasion.

## Introduction

During metastasis, cells must degrade the surrounding extracellular matrix (ECM) to create a path away from the solid tumor. Matrix metalloproteases (MMPs) are a family of soluble or membrane-bound enzymes that cleave components of the ECM. MT1-MMP, also called MMP14, is a transmembrane domain-containing protein that plays a key role in regulating metastasis. MT1-MMP itself can cleave numerous ECM substrates, including collagen, fibronectin, laminin and gelatin (Ohuchi et al., 1997), and can also process and activate other MMPs, including MMP-2 (Sato et al., 1994) and MMP-13 (Knäuper et al., 1996), further enhancing its ability to drive invasion.

ECM degradation by cells is promoted by the formation of podosomes (dendritic cells, macrophages, osteoclasts) or invadopodia (cancer cells) – actin-rich membrane protrusions that act as the focal point for MMP-mediated ECM cleavage. Following biosynthesis and traffic to the plasma membrane, MT1-MMP is efficiently internalized by clathrin-mediated and caveolar endocytosis (Poincloux et al., 2009) before being trafficked to late endosomes where it accumulates. Endosomes provide a reservoir of MT1-MMP that ensure focal delivery of MT1-MMP to the invadopodia (Marchesin et al., 2015). Dysregulation of endosomal trafficking machineries, including the retrograde components SNX27 and retromer (Sharma et al., 2019), the ESCRT-0 component Hrs (MacDonald et al., 2018), or the endosomal Arp2/3 activator Wiskott-Aldrich syndrome protein and Scar homolog (WASH) (Monteiro et al., 2013; MacDonald et al., 2018; Marchesin et al., 2015) all impair MT1-MMP delivery to the cell surface, and subsequently impair ECM degradation.

Endosomes are dynamic intracellular organelles that play crucial roles in protein sorting within cells. Cargos at the limiting (outer) membrane of endosomes can be redistributed, and endosomes are subject to extensive remodeling. Cytosolic tubules or vesicles transfer content to and from endosomes, and vesicles can bud into the lumen of the endosome. These internal vesicles, known as intraluminal vesicles (ILVs), are formed through ESCRT-dependent or ESCRT-independent mechanisms (Babst, 2011; Trajkovic et al., 2008; van Niel et al., 2011; Baietti et al., 2012; Edgar et al., 2014) and endosomes containing ILVs are termed multivesicular endosomes/bodies (MVE/MVBs). MVEs can fuse with lysosomes to form endolysosomes - catalytic organelles where substrates are degraded (Bright et al., 2016). Alternatively, MVEs can fuse with the plasma membrane, where their released ILVs become termed ‘exosomes’ (Raposo et al., 1996).

A proportion of MT1-MMP recycles from late endosomes to invadopodia through the formation of endosomal tubules (Marchesin et al., 2015). Interestingly, exogenously expressed tagged-MT1-MMP rather localizes to ILVs of MVEs and not to the limiting membrane of endosomes (Rossé et al., 2014; Wenzel et al., 2024). Consistent with MT1-MMP localizing of ILVs, MT1-MMP has also been found on exosomes (Hakulinen et al., 2008; Wenzel et al., 2024). Indeed, MT1-MMP and other MMPs have been found on other types of extracellular vesicles (Shimoda and Khokha, 2017; Thuault et al., 2022). MT1-MMP positive extracellular vesicles maintain the ability to both cleave collagen and convert pro-MMP-2 to active MMP-2 (Hakulinen et al., 2008). However, the relationship between MT1-MMP on ILVs/exosomes and invadopodia is not well understood.

Invadopodia act as critical docking and secretion sites for exosomes (Hoshino et al., 2013) and impairing invadopodia formation reduces exosome release from cells. Similarly, exosomes appear to promote the formation and maintenance of invadopodia through a positive feedback loop (Hoshino et al., 2013).

Exosomes and extracellular vesicles can shape the tumor microenvironment. Once exosomes are released from a cell, they can be transferred to other cells, or can interact with the ECM to remodel its structure (Karampoga et al., 2022). Whilst many studies focus on exosomes being released away from cells, exosomes can also remain associated to their cell of origin. We previously demonstrated that expression of the anti-viral restriction factor, tetherin, retains exosomes at the surface of cells (Edgar et al., 2016). Tetherin plays an analogous role in the tethering of enveloped virions (Neil et al., 2008; Kaletsky et al., 2009; Mandana et al., 2009; Stewart et al., 2023) and midbody remnants (Presle et al., 2021). Tetherin is a type II integral membrane protein, that forms homodimers through conserved cysteine residues within its extracellular domain (Ohtomo et al., 1999). Tetherin expression correlates with poor survival in invasive breast cancer patients (Mahauad-Fernandez et al., 2018) and promotes the invasiveness of cancer cells (Mahauad-Fernandez et al., 2014), although the mechanism linking tetherin and invasion is not well understood.

In this study, we aimed to determine how endogenous MT1-MMP is trafficked to EVs, and to analyse whether tetherin expression regulates the retention and release of MT1-MMP positive EVs. We found that endogenous MT1-MMP localizes predominately to ILVs, rather than to the limiting membrane of endosomes, and that MT1-MMP is retained at the surface of cells by tethered exosomes. Loss of tetherin enhances MT1-MMP positive exosome release and reduces the ability of cells to degrade the ECM. Together, these data highlight tethered exosomes as a contributor to ECM degradation, and detail how tetherin expression promotes invasiveness by cancer cells.

## Results

### MDA-MB-231 cells release sEVs/exosomes bearing MT1-MMP

MDA-MB-231 cells are triple negative breast cancers that form invadopodia and express endogenous MT1-MMP (Chen et al., 1994; Monteiro et al., 2013; Sharma et al., 2019; Pedersen et al., 2020). To elucidate the endosomal trafficking of endogenous MT1-MMP we performed confocal microscopy of MDA-MB-231 cells stained for endogenous MT1-MMP and the following endosomal markers: EEA-1 for early endosomes, CD63 for multivesicular and late endosomes, Rab7 for late endosomes, LBPA for late endosomes and endolysosomes, and LAMP-1 for late endosomes, endolysosomes and lysosomes (Fig. 1A). Co-localization analysis showed that MT1-MMP localizes throughout the endosomal pathway (from early endosomes to lysosomes), but is enriched in multivesicular bodies and late endosomes, as suggested by higher levels of colocalization with CD63 and Rab7 (Fig. 1A and 1B) and confirming previous reports (Williams and Coppolino, 2011; Planchon et al., 2018; Pedersen et al., 2020). We also observed co-localization of MT1-MMP with the retromer complex subunit VPS35 and the sorting nexin SNX27 that have been previously shown to facilitate the recycling of MT1-MMP from endosomes to the plasma membrane (Sharma et al., 2019) (Fig. S1A). Nevertheless, while MT1-MMP positive endosomes mostly appeared as “filled circles”, VPS35 and SNX27 staining appeared as “empty circles” (“donut shape”). Line scan analysis of endosomes positive for MT1-MMP and VPS35/SNX27 confirmed that VPS35 and SNX27 mostly decorate the outer profile of endosomes, while MT1-MMP appears enriched in the lumen of such endosomes (Fig. S1B).

**Figure 1.**
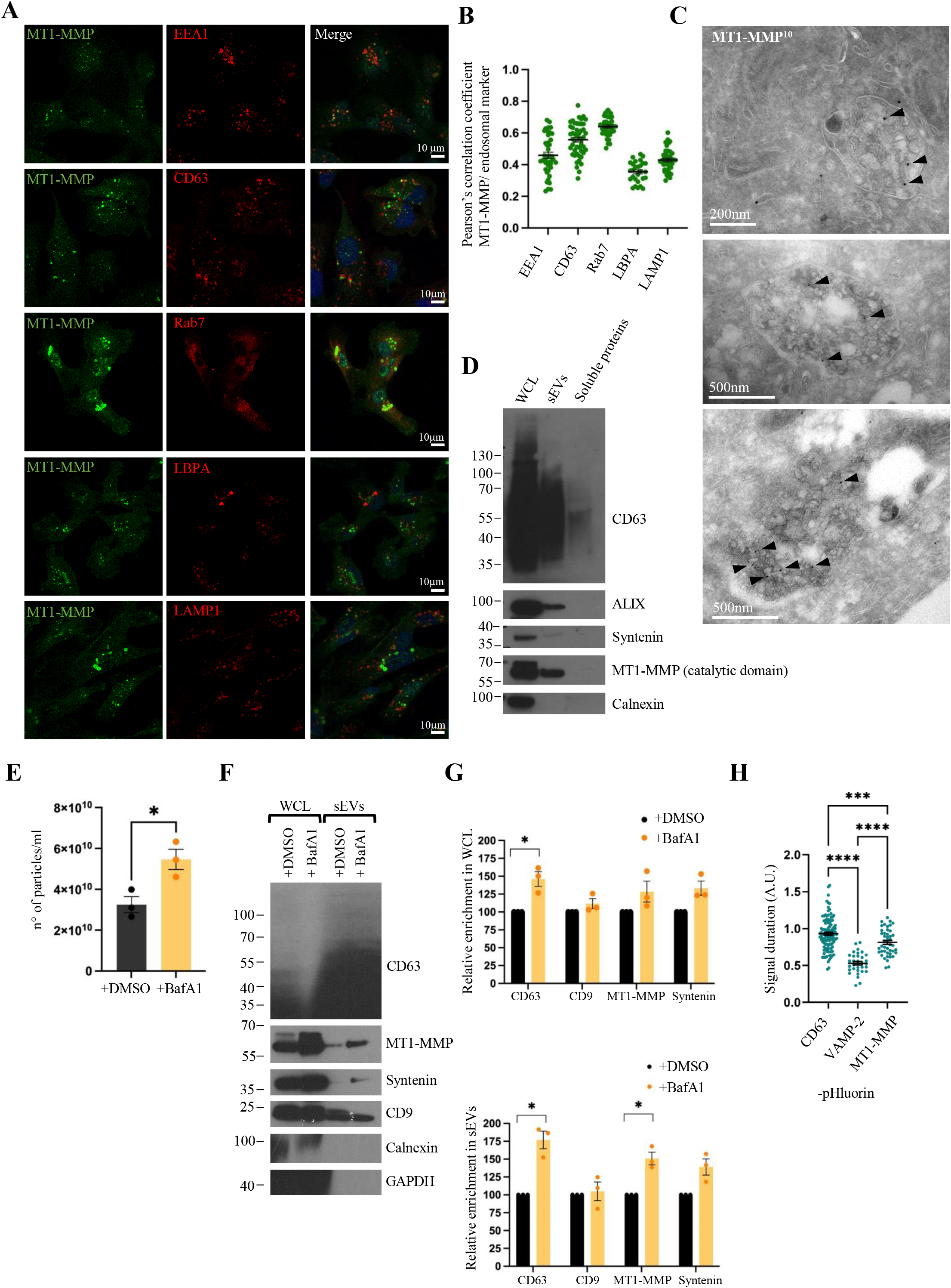
MDA-MB-231 cells release sEVs/exosomes bearing MT1-MMP. **(A)** Confocal analysis of endogenous MT1-MMP localization with different endosome/lysosome markers in MDA-MB-231 WT cells. Scale bars = 10 μm. **(B)** Pearson’s correlation coefficient (mean ± SEM) of MT1-MMP and endosome/lysosome markers. n ≥ 28 cells from 3 independent experiments. **(C)** Immuno EM micrographs of MDA-MB-231 WT cells labelled with MT1-MMP and PAG 10 nm. Scale bar = 200 nm and 500 nm. **(D)** Western blotting analysis of whole cell lysates (WCL), sEV enriched fraction, and soluble protein enriched fraction. **(E)** Concentration (mean ± SEM) of sEVs isolated from MDA-MB-231 WT cells treated with DMSO or BafA1 for 16 hours and analyzed by NanoFCM. n = 3 biologically independent experiments, unpaired *t* test, p= 0.0249. **(F)** Western blotting analysis of whole cell lysates (WCL) and sEVs from MDA-MB-231 cells treated with DMSO or BafA1. **(G)** Relative enrichment (mean ± SEM) of the protein content in WCL or sEVs from MDA-MB-231 cells treated with DMSO or BafA1. n = 3 biologically independent experiments, unpaired *t* test with Welch correction, * p= 0.046 for CD63 WCL, * p=0.024 for CD63 EVs, * p=0.028 for MT1-MMP EVs. **(H)** Comparison (mean ± SEM) between signal duration of fusion events of CD63-, MT1-MMP-, and VAMP2- pHluorin. n= 109 fusion events for CD63, n= 32 for VAMP-2, n=44 for MT1-MMP, from 3 biologically independent experiments, Ordinary one-way ANOVA, *** p= 0.0009, **** p<0.0001. All molecular weights are in kDa.

Cryo immunogold transmission electron microscopy (TEM) provides sufficient resolution to determine antigen localization between subdomains of endosomes, including differentiating labelling to the outer membrane and ILVs, so we employed this method to further determine the sub-organellar localization of MT1-MMP. Immunogold TEM revealed a significant enrichment of MT1-MMP on ILVs of multivesicular endosomes (MVEs) (Fig. 1C). Cargos trafficked to ILVs are degraded if the MVE fuses with the lysosome, or alternatively MVEs can fuse with the plasma membrane and release their ILVs in the extracellular space, where secreted ILVs are termed ‘exosomes’ (Raposo et al., 1996). To assess the capacity of MT1-MMP to undergo lysosomal degradation, as also suggested by the colocalization of MT1- MMP with LBPA and LAMP1 (Fig. 1, A and B), we treated cells with the lysosomal protease inhibitor leupeptin for 2 or 4 hours. Western blot analysis of cell lysates showed a small increase in MT1-MMP levels in leupeptin treated cells (Fig. S1, C and D). Confocal microscopy confirmed the accumulation of MT1-MMP in LAMP-1 positive compartments upon treatment with leupeptin (Fig. S1E). A similar accumulation was observed when cells were treated with Bafilomycin A1 (BafA1), an inhibitor of the V-ATPase (required for lysosomal acidification and degradative function) (Fig. S1F). Overall, these observations demonstrate that, at least a fraction of MT1-MMP is trafficked to lysosomes where it undergoes lysosomal degradation.

We also stained cells with MagicRed, a live fluorescent reporter for Cathepsin B activity, that can be used as a marker of degradative endolysosomes (Bright et al., 2016). Similarly to CD63, MT1-MMP was found to only partially colocalize with MagicRed, confirming the presence of MT1-MMP in endosomes with degradative capacity (Fig. S1, G and H). MT1-MMP was also observed in MagicRed negative (non-catalytic) organelles, supporting the idea that a fraction of MT1-MMP might escape lysosomal degradation and be fated for exosome release.

To assess the capacity of MDA-MB-231 to release endogenous MT1-MMP on EVs/exosomes, as previously described (Hakulinen et al., 2008; Hoshino et al., 2013; Beghein et al., 2018), we isolated small EVs (sEVs) using size exclusion chromatography (SEC), allowing the “EV enriched fraction” to be separated from the “soluble protein enriched fraction”. Transmission electron microscopy (Fig. S1I) and Flow Nano Analyzer (NanoFCM) analysis (Fig. S1J) confirmed the presence of sEVs in the “EV enriched fraction”. Western blot analysis of the two fractions confirmed the presence of sEVs markers (CD63, ALIX (ALG-2-interacting Protein X), syntenin) and the absence of the ER marker calnexin in the “EV enriched fraction” (Fig. 1D). We also identified the full-length form of MT1-MMP in the sEV enriched fraction and its absence in the soluble protein enriched fraction (Fig. 1D), confirming that a catalytically active transmembrane form of MT1-MMP is released through sEVs.

Treating cells with Bafilomycin A1 (BafA1) specifically increases the release of exosomes (derived from the endosomal system) (Edgar et al., 2016; Mathieu et al., 2021). Electron microscopy of BafA1 treated MDA-MB-231 cells showed the presence of enlarged endolysosomes (Fig. S1K), consistent with our immunofluorescence microscopy observations (Fig. S1F). NanoFCM analysis confirmed that BafA1 treatment induced an increase of exosome release in MDA-MB-231 cells (Fig. 1E). Western blot analysis demonstrated that CD63, syntenin and MT1-MMP levels were increased in sEV fractions upon BafA1 treatment (Fig. 1F and G), suggesting that MT1-MMP positive sEVs originate in the endosomal system (exosomes). In contrast, CD9 levels in sEVs were not affected, suggesting that CD9 positive vesicles might originate mostly from the plasma membrane (Fig. 1F and G), in agreement with recent work (Mathieu et al., 2021).

Total internal reflection fluorescence (TIRF) microscopy of live cells expressing CD63 tagged with a pH-sensitive fluorescent protein (pHluorin) has been previously used to quantify the dynamics of MVEs - PM fusion (hence exosome release) (Verweij et al., 2018). As we observed the presence of MT1-MMP on sEVs/exosomes we decided to compare the dynamics of CD63-pHluorin release to those of MT1-MMP-pHluorin (Figure 1H and video 1). Additionally, we compared them with the dynamics of VAMP-2-pHluorin, a SNARE protein that inserts in the plasma membrane upon fusion of endosomes with the plasma membrane. We observed that the signal duration of both CD63- and MT1-MMP – pHluorin were significantly longer than that of VAMP-2-pHluorin, suggesting that, as for CD63, the secretion of MT1-MMP-pHluorin corresponds to an event of exosome release (Figure 1H and video 1).

Overall, these observations showed that MT1-MMP is enriched in ILVs of MVEs and can be released in the extracellular environment through exosomes.

### MDA-MB-231 cells express tetherin and tether MT1-MMP-exosomes to the cell surface

Exosomes that are released in the extracellular space by donor cells can diffuse and be taken up by distant recipient cells. Alternatively, exosomes can remain attached to the surface of donor cells through the action of tetherin (Edgar et al., 2016). These vesicles appear as clusters of exosomes on the surface of donor cells.

As breast tumors and triple-negative human breast cancer cell line such as MDA-MB-231 have been previously shown to express high levels of tetherin (Mahauad-Fernandez et al., 2014), we decided to investigate the capacity of these cells to tether exosomes. Western blot analysis of MDA-MB-231 cell lysates confirmed that these cells express high levels of tetherin and tetherin was also found on exosomes isolated by SEC (Fig. 2A). Immunofluorescence microscopy demonstrated that tetherin localized abundantly to late endosomes (Fig. S2, A and B). Moreover, tetherin colocalized strongly with MT1-MMP in punctate organelles (Fig. 2, B and C), consistent with endosomal staining (Fig. 1, A and B). Similarly to MT1-MMP, tetherin accumulated intracellularly upon treatment with BafA1 (Fig. 2, D and E), specifically in LAMP-1 positive compartments (Fig. S2C). The levels of tetherin in EV-enriched fractions were also increased upon BafA1 treatment, suggesting that tetherin positive EVs are exosomes (Fig. 2, D and E). Immunofluorescence staining of non-permeabilized cells (surface staining) showed the presence of tetherin on the cell surface, which became less uniform upon treatment of cells with BafA1 (Fig. S2D).

**Figure 2.**
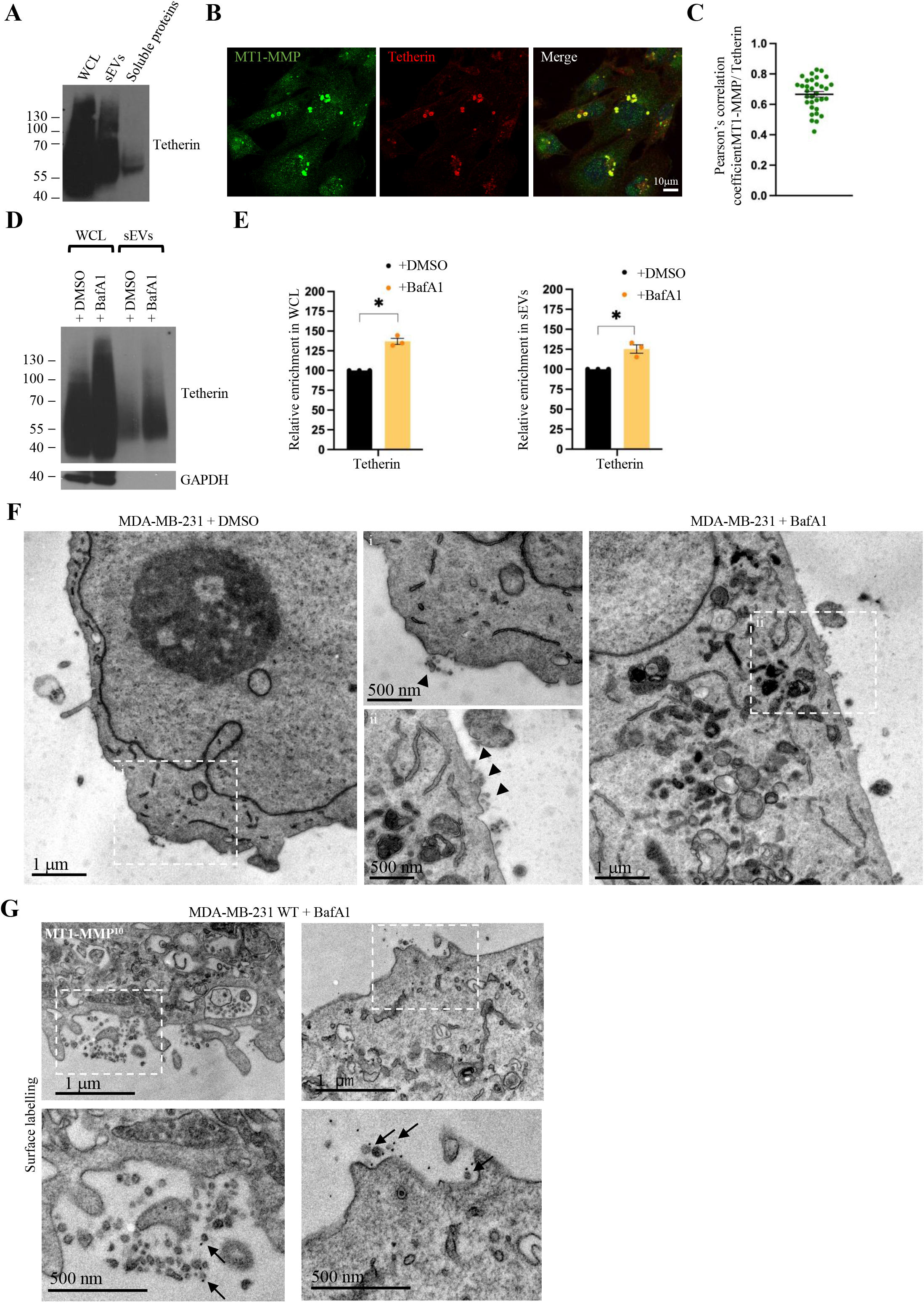
MDA-MB-231 express tetherin and tether MT1-MMP- exosomes at the cell surface. **(A)** Western blotting analysis of WCL, sEV enriched fraction and soluble protein enriched fraction. **(B)** Confocal analysis of endogenous MT1-MMP localization with Tetherin. Scale bar = 10 μm. **(C)** Pearson’s correlation coefficient (mean ± SEM) of MT1-MMP and Tetherin. n= 33 cells from 2 biologically independent experiments. **(D)** Western blotting analysis of WCL and sEVs from MDA-MB-231 cells treated with DMSO or BafA1 for 16 hours. **(E)** Relative enrichment (mean ± SEM) of tetherin content in WCL or sEVs from MDA-MB-231 cells treated with DMSO or BafA1. n = 3 biologically independent experiments, unpaired *t* test with Welch correction, * p= 0.010 for WCL, * p= 0.041 for EVs. **(F)** EM micrograph of MDA-MB-231 WT cells treated with DMSO or BafA1. Magnification shows the presence of tethered EVs (arrowheads) near the plasma membrane. Scale bars = 1 μm and 500 nm. **(G)** EM micrograph of MDA-MB-231 WT cells treated with BafA1 for 16 hours and labelled for MT1-MMP (PAG10). Arrows indicate MT1-MMP positive vesicles tethered at the cell surface. Scale bars = 1 μm and 500 nm.

However, as confocal microscopy does not allow one to definitively observe tethered EVs at the cell surface of cells, we performed conventional electron microscopy (Fig. 2F). Only a few small EVs were observed in proximity of the cell surface of control cells, hence, to better analyze MDA-MB-231 cells, we induced exosome release using BafA1 as this increased the frequency of tethered exosome clusters at the cell surface (Fig. 2F). Surface labelling TEM was performed using an anti-MT1-MMP antibody and revealed the presence of MT1-MMP (Fig. 2G) on tethered exosomes.

Overall, we conclude that MDA-MB-231 cells express high levels of tetherin and can tether exosomes bearing MT1-MMP at their surface.

### Tetherin expression modulates release of MT1-MMP/exosomes and their presentation at the surface of cells

To further investigate the role of tetherin in MDA-MB-231 cells and its relationship with MT1-MMP and exosomes, we generated a Bst-2/tetherin knock out (Bst-2 KO) cell line by transducing Bst-2 gRNAs into a Cas9 stable cell line, that was also used for our analysis. The loss of tetherin protein in Bst-2 KO cells was demonstrated by Western blot, immunofluorescence microscopy and flow cytometry (Fig. 3, A-C).

**Figure 3.**
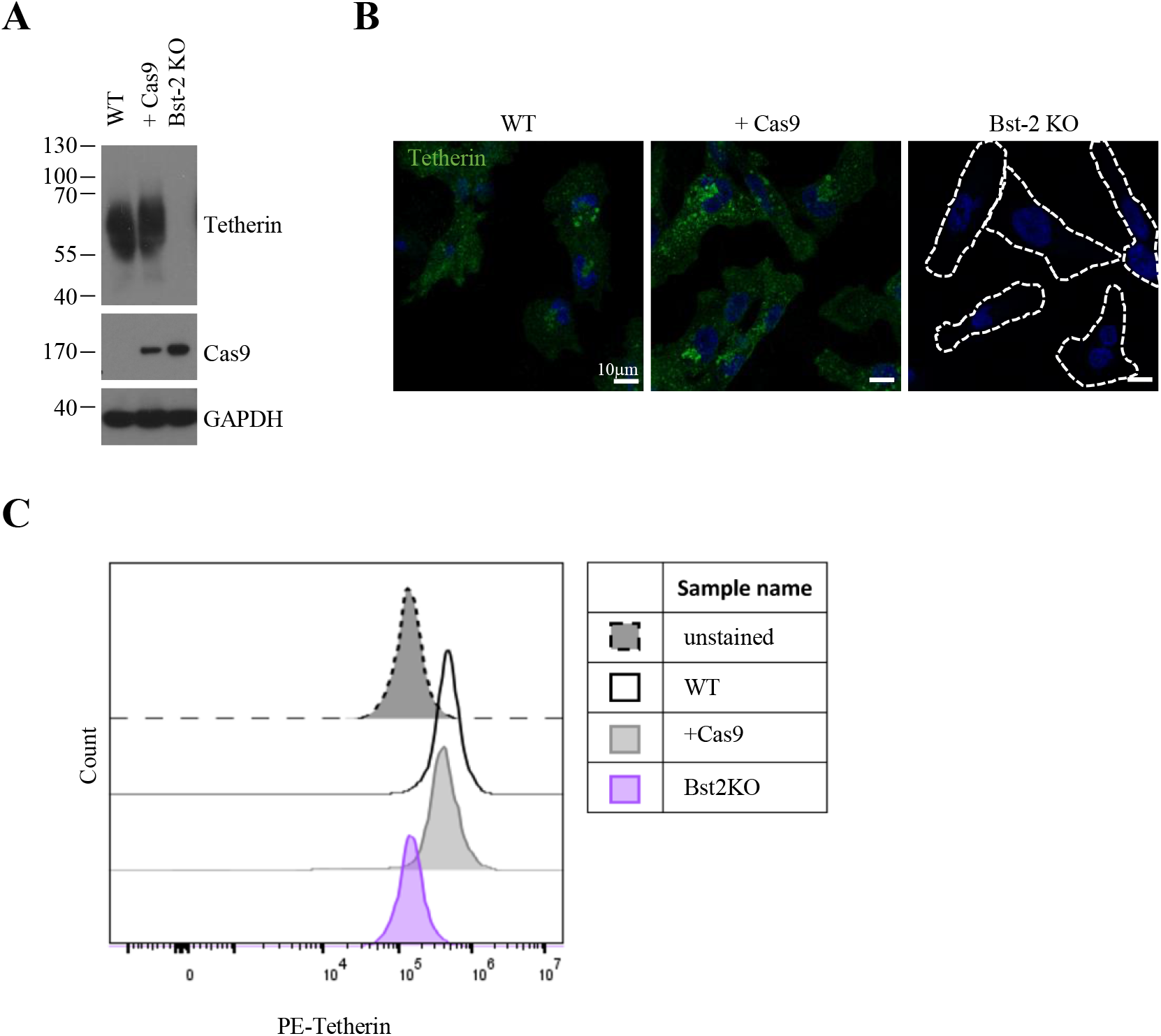
Characterization of MDA Bst-2 cell lines. **(A)** Western blotting analysis of tetherin and Cas9 expression in MDA-MB-231 cell lines. **(B)** Confocal analysis of MDA-MB-231 cell lines stained for tetherin. Scale bars = 10 μm. **(C)** Flow cytometry analysis of surface tetherin levels in MDA-MB-231 cell lines.

As tetherin has been previously shown to interact with MT1-MMP and potentially affect its trafficking (Gu et al., 2012; Fan et al., 2016), we investigated if the endosomal localization of MT1-MMP was affected by tetherin loss. Confocal microscopy and co-localization analysis of MT1-MMP with endosomal markers did not show any major change in MT1-MMP localization in Bst-2 KO cells (Fig. 4, A and B, Fig. S3A and S3B). Colocalization of MT1-MMP with the retromer complex subunit VPS35 was not affected (Fig. S3C), suggesting that tetherin expression similarly did not influence the retromer/SNX27-mediated recycling of MT1-MMP.

**Figure 4.**
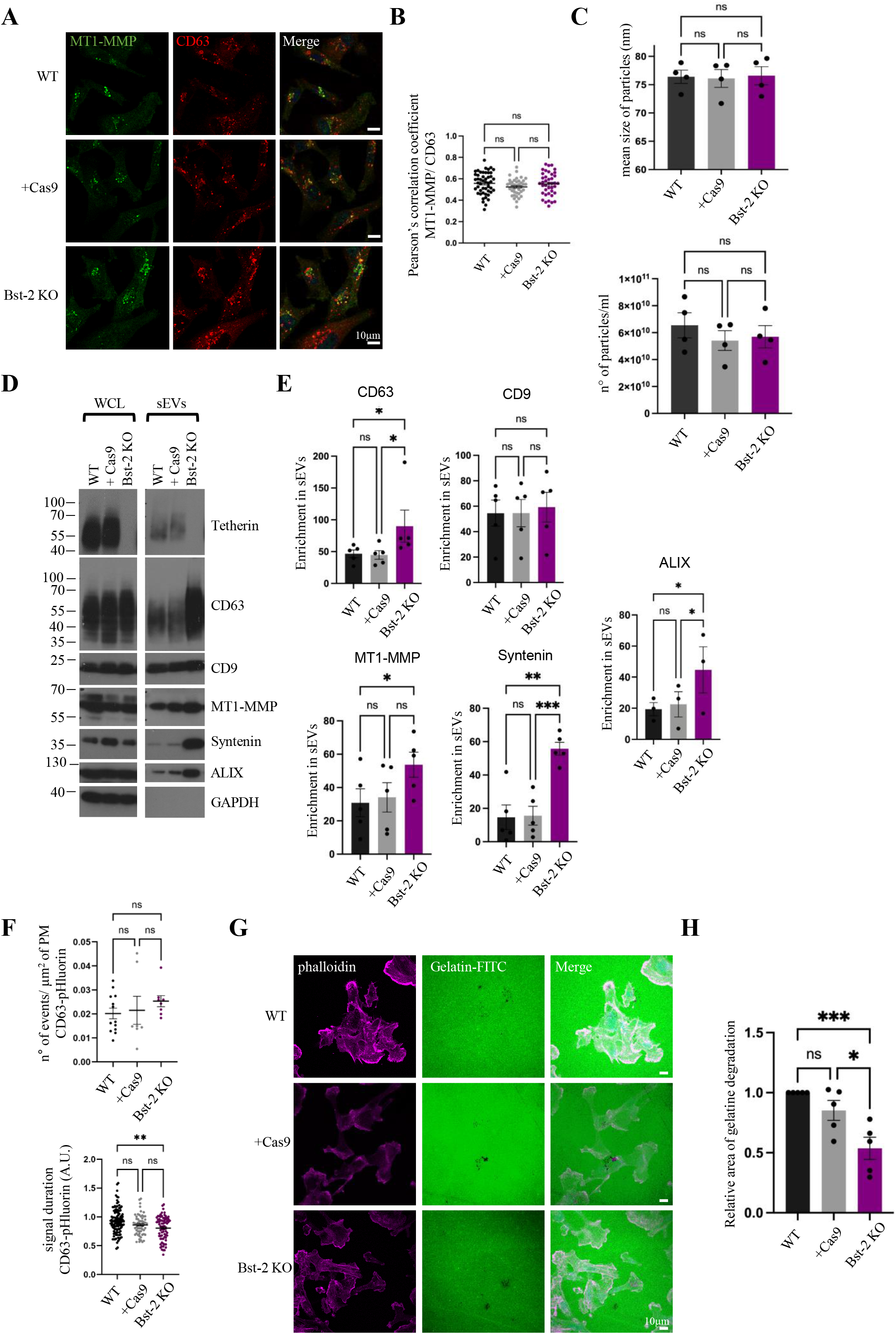
Bst-2 expression modulates release of MT1-MMP/exosomes and their presentation at the surface of cells. **(A)** Confocal analysis of endogenous MT1-MMP localization with CD63 in MDA-MB- 231 cell lines. Scale bars = 10 μm. (**B**) Pearson’s correlation coefficient of MT1-MMP and CD63. n = 49 for WT, 39 for Cas9, 40 for Bst-2 KO, cells from 3 biologically independent experiments, mean ± SEM, ordinary one-way ANOVA. (**C**) Size and concentration (mean ± SEM) of sEVs isolated from MDA-MB-231 cell lines and analyzed by NanoFCM. n = 4 biologically independent experiments, Ordinary one- way ANOVA. (**D**) Western blotting analysis of WCL and sEV from MDA-MB-231 cell lines. **(E)** Enrichment of the protein content (mean ± SEM) in sEVs from MDA-MB- 231 cell lines. n = 5 biologically independent experiments, Friedman test for CD63, * p=0.0114 and * p=0.0269; one-way ANOVA for MT1-MMP, CD9, syntenin, and ALIX, * p=0.0326 for MT1-MMP, ** p=0.0021 and *** p=0.0007 for syntenin, * p=0.0298 and * p=0.0444 for ALIX. **(F)** Fusion activity of CD63-pHluorin. n = 13 for WT, 7 for Cas9, 8 for Bst-2 KO, cells from 3 biologically independent experiments, Welch ANOVA test. Signal duration of fusion events of CD63-pHluorin. n = 109 for WT, 61 for Cas9, 76 for Bst-2 KO, fusion events from 3 independent experiments, Kruskal- Wallis test, ** p=0.0031. **(G)** Confocal analysis of gelatin degradation assay. Cells were plated on gelatin-FITC coated coverslips, incubated for 5 hours, fixed, and stained with phalloidin. Scale bars = 10 μm. **(H)** Quantification of the relative area of degraded gelatin (mean ± SEM). n = 5 biologically independent experiments, Ordinary one-way ANOVA, * p=0.0121 and *** p=0.0007.

Tetherin expression can modulate the balance of extracellular vesicle release versus retention, including exosomes (Edgar et al., 2016), midbody remnants (Presle et al., 2021) and enveloped viruses (Neil et al., 2008). sEVs/ exosomes were isolated from the different cell lines using SEC. NanoFCM analysis of isolated EVs revealed no difference in EV size or EV number (Fig. 4C). However, Western blots analysis of the same samples showed that tetherin loss significantly enhanced the levels of CD63, ALIX, syntenin and MT1-MMP, but not CD9, in EV fractions (Fig. 4D and E), while cellular levels of each marker remained unchanged (Fig. S3D). These data confirm that loss of tetherin impacts EV release of at least some EV subpopulations. As sEVs isolated from the cell culture media represent a heterogenous mix of vesicles with different origin whose release might or might not be affected by tetherin expression, we decided to specifically study the release of endosome derived exosomes using CD63-pHluorin and of MT1-MMP positive exosomes using MT1- MMP-pHluorin (Fig. 4F and S3E; video 2 and 3).

While the number of MVE-PM fusion events (identified by CD63 or MT1-MMP- pHlourin activity) was not affected by tetherin expression, the duration of the fusion events was decreased in Bst-2 KO cells. This further confirms that in absence of tetherin, exosomes do not stay in proximity of the cell surface and diffuse more rapidly in the culture media (Fig. 4F and S3E; video 2 and 3).

Exposure of MT1-MMP on the cell surface is required to allow cancer cells to degrade the ECM in the pericellular space and therefore invade and migrate in the interstitial space. As tetherin expression has been previously associated with increased migratory capacity of breast cancer cells (Yi et al., 2013; Mahauad-Fernandez et al., 2014; Naushad et al., 2017; Mahauad-Fernandez et al., 2018), we decided to investigate the impact of tetherin loss on the capacity of MDA-MB-231 cells to degrade the ECM and form invadopodia. Cells were seeded on coverslips coated with fluorescent gelatin and incubated for 5 hours to allow gelatin degradation (Fig. 4, G and H). Bst-2 KO cells were less able to degrade the gelatin, when compared to WT or +Cas9 cells. Overall, these data confirm that tetherin loss enhances the release of exosomes, and MT1-MMP from cells, which impairs the capacity of cells to remodel the local ECM.

### Tetherin regulates invadopodia formation and ECM degradation

To ensure that the impact of tetherin loss on ECM degradation was direct, and not caused by off-target effects, we generated a tetherin rescue cell line. Tetherin-HA was stably expressed in Bst-2 KO MDA-MB-231 cells and expression was confirmed by Western blot and immunofluorescence microscopy (Fig. 5, A and B). The ability of cells to tether exosomes was examined by TEM (Fig. 5C) and the frequency of sEVs calculated (Fig. 5D). The frequency of sEVs at the surface of cells significantly decreased upon tetherin loss and was restored in rescue cells.

**Figure 5.**
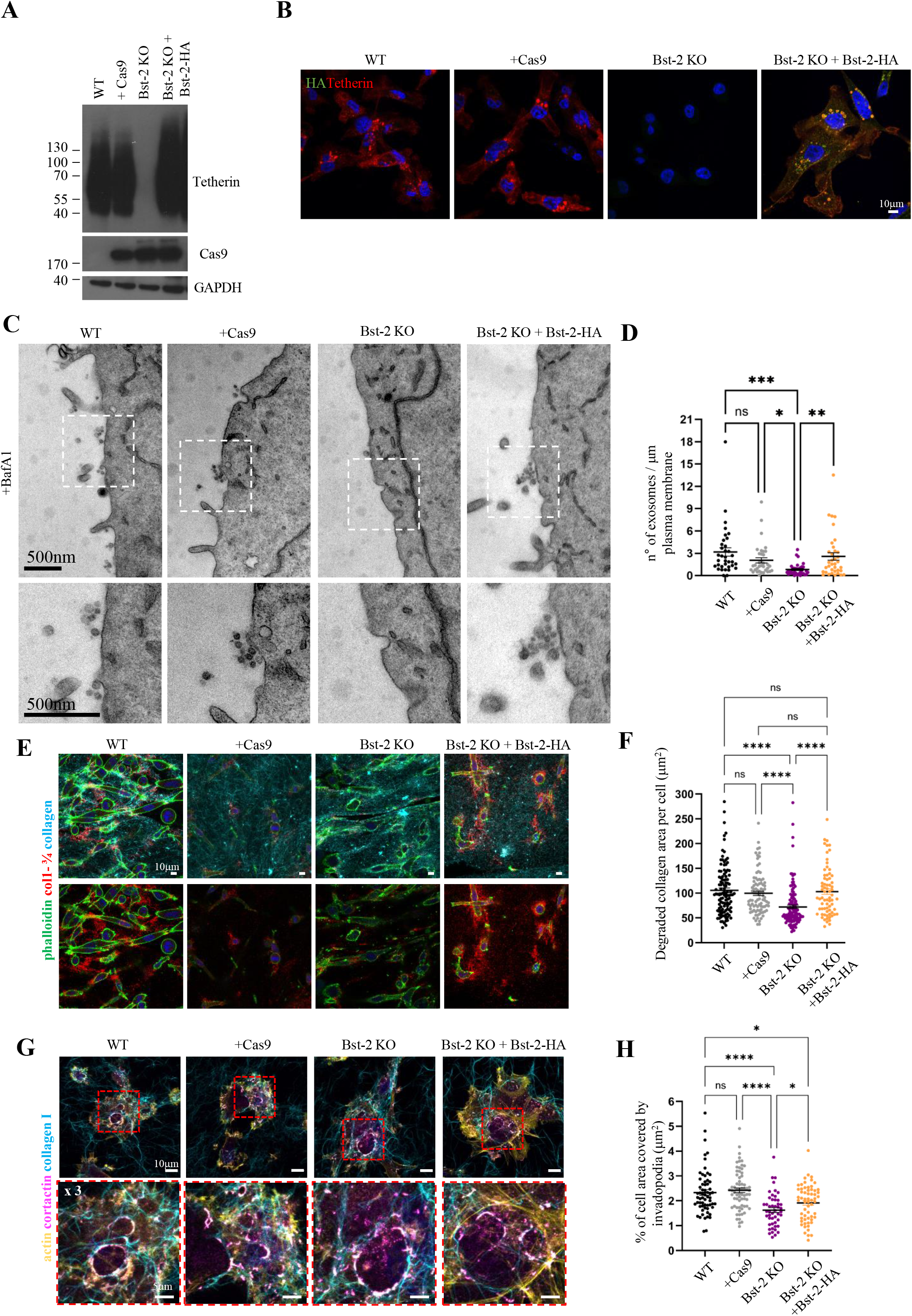
Bst-2 regulates ECM degradation and invadopodia formation. **(A)** Western blotting analysis of tetherin expression in MDA-MB-231 cell lines. **(B)** Confocal analysis of MDA-MB-231 cell lines stained for tetherin and HA. Scale bars = 10 μm. **(C)** EM micrograph of MDA-MB-231 cell lines treated with BafA1. Magnification shows the presence of tethered EVs near the plasma membrane. Scale bars = 500 nm. **(D)** Quantification (mean ± SEM) of the number of tethered sEVs along the plasma membrane (number of sEVs/μm of plasma membrane). n = 35 cells from 3 biologically independent experiments, one-way ANOVA, * p= 0.0402, ** p=0.0034, *** p=0.0001. **(E)** Confocal analysis of collagen degradation assay. MDA-MB-231 cells were embedded in 3D fluorescent collagen gels, incubated for 24 hours, fixed, and stained with phalloidin and for the cleaved collagen new epitope using anti-Col-3/4 antibody. Scale bars = 10 μm. **(F)** Quantification of pericellular collagen degradation (mean ± SEM) expressed as average degraded collagen area per cell. n = 112 for WT, 92 for Cas9, 126 for Bst-2 KO, 71 for Bst-2 KO+ Bst-2-HA, confocal images (≥ 400 cells) from at least 3 biologically independent experiments, Kruskal-Wallis test, **** p<0.0001. **(G)** Confocal analysis of invadopodia formation assay. Cells were plated on a thin layer of fluorescent collagen, incubated for 60 minutes, fixed, and stained with cortactin and phalloidin. Magnification shows the presence of linear invadopodia along collagen fibers. **(H)** Quantification of the cell area covered by invadopodia structures (mean ± SEM). n = 62 for WT, 68 for Cas9, 48 for Bst-2 KO, 60 for Bst-2 KO + Bst-2-HA, cells from at least 3 biologically independent experiments, Kruskal-Wallis test, * p=0.0271, * p=0.0225, **** p<0.0001.

To evaluate the effect of tetherin expression on ECM degradation, we conducted assays in 3D cultures. Cells were embedded in 3D fluorescent collagen I gels and incubated for 24 hours to allow degradation of the collagen. Subsequently 3D collagen gels were stained with the antibody collagen ¾, which specifically recognize degraded collagen (antibody against an epitope that is exposed only upon cleavage of collagen by metalloproteinases) (Fig. 5, E and F). Similarly to the gelatin degradation assay (Fig. 4, F and G), we observed that Bst-2 KO cells were less able to degrade collagen, while tetherin rescue cells showed similar levels of collagen cleavage relative to WT cells (Fig. 5, E and F). A small increase in collagen cleavage was observed in tetherin rescue cells, and this may be due to the slightly higher expression of ectopic tetherin in the rescue cells, relative to endogenous tetherin in WT cells.

Given tetherin expression can be induced by type I interferons (IFNs) (Ohtomo et al., 1999; Neil et al., 2008), we examined whether IFN-mediated induction of tetherin could enhance collagen degradation. MDA-MB-231 WT cells were treated with IFN-α for 16 hours and we confirmed that tetherin expression was increased by immunofluorescence microscopy and western blotting (Fig. S4, A and B). Collagen degradation assays were performed in mock or IFN-α treated MDA-MB-231 cells, and we observed that IFN-α treated cells were able to degrade collagen similarly - showing a mildly elevated but not significant increase in collagen cleavage (Fig. S4, C and D).

As the capacity to degrade the extracellular matrix is closely associated to the capacity of cancer cells to form invadopodia, we assessed the ability of MDA-MB- 231 cell lines to form linear invadopodia along collagen fibers (Fig. 5, G and H). Cells were seeded on a thin layer of fluorescent collagen, incubated for 1 hour to allow invadopodia formation, and stained for invadopodia markers cortactin and actin (Fig. 5G). When the area of each cell that was covered by invadopodia was quantified, we found that Bst-2 KO cells had less invadopodia than WT cells, while rescue cells demonstrated a significant restoration in invadopodia formation relative to KO cells (Fig. 5H).

Together, these data demonstrate that tetherin expression modulates the relative retention and release of MT1-MMP positive exosomes from MDA-MB-231 cells, and that the levels of released MT1-MMP correlate with the capacity of cells to degrade the pericellular ECM.

## Discussion

Extracellular matrix degradation depends on the delivery of MT1-MMP to the extracellular space surrounding migrating cancer cells. In this study, we demonstrate that in addition to being delivered to the plasma membrane, MT1-MMP is also present on small extracellular vesicles (sEVs)/exosomes, which are anchored to the surface of migrating cells via tetherin. The loss of tetherin from cells reduces the abundance of MT1-MMP-positive exosomes at the surface of cells, and exosomes are instead released to the extracellular environment, with a concomitant decrease in the invasiveness of cells. This study is the first to show that tethered exosomes play a functional role in cancer pathogenesis.

Previous studies have identified Rab7-positive late endosomes as the primary storage site for MT1-MMP (Williams and Coppolino, 2011; Planchon et al., 2018; Pedersen et al., 2020), though much of data relies on the use of ectopically expressed forms of MT1-MMP (Chevalier et al., 2016; Rossé et al., 2014; Wenzel et al., 2024). Here, we extend these findings by localizing endogenous MT1-MMP in MDA-MB-231 cells predominately to Rab7-positive late endosomes and CD63-positive multivesicular endosomes / lysosomes (Fig. 1, A and B). Using cryo immunogold electron microscopy we demonstrate that MT1-MMP is localized to ILVs, rather than the limiting membrane or tubules of endosomes (Fig. 1C). This traffic to ILVs is likely to preclude MT1-MMPs availability for traffic to invadopodia via endosomal tubules. Fusion of CD63 positive compartments with the plasma membrane releases exosomes (Verweij et al., 2018) and we confirm that MT1-MMP-positive extracellular vesicles (Fig. 1D-G) are exosomes that originate in the endosomal system (Fig. 1H).

Several pieces of evidence indicate that the balance of MT1-MMP traffic to ILVs/exosomes versus the limiting endosomal membrane/plasma membrane is an important determinant of MT1-MMP function (Thuault et al., 2022), although the precise molecular mechanisms governing MT1-MMP sorting to ILVs and exosomes remain unclear and warrant further investigation.

Mono-ubiquitination of MT1-MMP at Lys^581^ alters MT1-MMP trafficking, enhancing its presence at the surface of cells and promoting cell invasion (Eisenach et al., 2012). Ubiquitinated endosomal cargoes can be recognized by the endosomal sorting complex required for transport (ESCRT) machinery and are sorted into ILVs (Raiborg and Stenmark, 2009). ILVs are degraded if the MVE fuses with the lysosome, or become exosomes if the MVE fuses with the plasma membrane. ILVs can be generated through at least two distinct molecular mechanisms (Stuffers et al., 2009; van Niel et al., 2011; Baietti et al., 2012; Edgar et al., 2014) – namely ESCRT-dependent and ESCRT-independent pathways. Despite the ubiquitination of MT1- MMP, it seems likely that it is trafficked to ILVs in an ESCRT-independent manner. Knockdown of the ESCRT-0 component, Hrs, impacts exosome release but does not affect the presence of MT1-MMP on exosomes (Hoshino et al., 2013). Rather, Hrs may help regulate endosomal recycling of MT1-MMP via the actin nucleating factor Wiscott-Aldrich syndrome protein and SCAR homologue (WASH) (MacDonald et al., 2018). Additionally, several studies have found MT1-MMP to interact with tetraspanins (Schröder et al., 2013; Lafleur et al., 2009) and tetraspanins regulate ESCRT-independent ILV formation (van Niel et al., 2011; Larios et al., 2020). The C-terminal PDZ-binding motif of MT1-MMP (DKV^582^) has been implicated in its intracellular traffic (Wang et al., 2004; Sharma et al., 2019) and given the PDZ-domain containing protein syntenin is central to a well characterized ILV/exosome biogenesis pathway (Baietti et al., 2012), its potential role in MT1-MMP sorting should be further investigated (Thuault et al., 2022).

The sorting of MT1-MMP to ILVs is likely necessary for its subsequent lysosomal degradation. Our data show that inhibiting lysosomal activity using leupeptin or BafA1 results in an accumulation of cellular MT1-MMP (Fig. S1C-F, Fig. 1, F and G), highlighting the importance of lysosomal degradation and acidification in maintaining MT1-MMP levels and activity, as previously reported (Maquoi et al., 2003). We also found the presence of MT1-MMP in non-degradative endosomal compartments (negative for the Cathepsin B substrate MagicRed) (Bright et al., 2016) (Fig. S1G), in agreement with previous observations (Planchon et al., 2018), and in compartments positive for CD63, a marker of secretory MVEs (Verweij et al., 2018, 2022), and of secretory endo-lysosome-related organelles (ELROs) (Delevoye et al., 2019). MVEs and ELROs share common effectors such as the Rab GTPases Rab27A and Rab27B that are required for their docking to the PM and secretion (Fukuda, 2013; Ostrowski et al., 2010) and interestingly, Rab27 knock down can downregulate MT1- MMP recycling (Macpherson et al., 2014), invadopodia formation and ECM degradation (Hoshino et al., 2013). Hence, our data highlight that MT1-MMP is stored in secretory and non-degradative late endosomes, likely in a pre- endolysosomal stage, from where it can be trafficked towards the plasma membrane.

Previous studies have observed and characterized the formation of tubules containing MT1-MMP emanating from late endosomes, that involve the motor adaptor proteins JIP3/4 and the WASH complex (Monteiro et al., 2013; Marchesin et al., 2015), likely in concert with the retromer complex and SNX27 (Sharma et al., 2019) or the ESCRT-0 subunit HRS (MacDonald et al., 2018), and the importance of these tubules for the delivery of MT1-MMP to invadopodia.

Supporting previous observations (Hoshino et al., 2013; Beghein et al., 2018), our combined microscopy analysis shows that endogenous MT1-MMP is trafficked to ILVs of MVBs (Fig. 1A, 1C, S1A and B) and suggest that an efficient way to deliver MT1-MMP to the pericellular space might be through fusion of MVEs with the PM and subsequent tethering of MT1-MMP-exosomes by tetherin (Fig. 2G). Several molecular machineries involved in the fusion of MVEs with the plasma membrane regulate MT1-MMP trafficking and delivery to invadopodia. This is the case of the aforementioned Rab27, the SNAREs VAMP7 (Steffen et al., 2008; Fader et al., 2009) and SNAP23 (Williams et al., 2014; Wei et al., 2017) or the recently identified role for Protudin-mediated ER-endosome contact sites that regulate both MT1-MMP exocytosis and invadopodia formation (Pedersen et al., 2020). Overall, it appears that several pathways regulating late endosomes motility, tubulation and fusion with the plasma membrane would co-operate to deliver MT1-MMP to the pericellular space, therefore contributing to invadopodia formation and ECM degradation.

Our data demonstrate that the invasiveness of cells correlates with the retention of MT1-MMP of tethered exosomes (Fig. 4, B-E). Expression of tetherin is required for the anchoring of exosomes to the plasma membrane of cells (Edgar et al., 2016) (Fig. 5, C and D). Tetherin is overexpressed in a number of cancers, including multiple myeloma (Rew et al., 2005), ovarian cancer (Yang et al., 2022), and breast cancer (Cai et al., 2009; Sayeed et al., 2013), where its high expression is linked to poor prognosis (Sayeed et al., 2013; Mahauad-Fernandez et al., 2018). Several studies have demonstrated that tetherin can promote cell adhesion (Mahauad-Fernandez et al., 2018; Yoo et al., 2011), migration (Liu et al., 2018) and metastasis (Cai et al., 2009; Mahauad-Fernandez et al., 2014, 2018). However, the molecular mechanism linking tetherin with metastasis is poorly defined.

Our findings show that tetherin plays a key role in regulating MT1-MMP-positive exosomes, enhancing their retention at the cell surface. When tetherin is lost, exosomes diffuse away from the cells (Figs. 4, A-E, 5, C and D), leading to an increased release of MT1-MMP from the cells (Fig. 4, B-E). Consistent with previous studies (Mahauad-Fernandez et al., 2018), we confirmed that tetherin loss reduces the ability of cancer cells to degrade the extracellular matrix (ECM) (Fig. 4, G and H, Fig. 5, E-H), and found that this loss promotes the release of MT1-MMP-containing exosomes into the pericellular space. Additionally, we observed that tetherin depletion decreases the presence of invadopodia along collagen fibres (Fig. 5, G and H). Invadopodia formation and maturation involve multiple stages: precursor formation, protease acquisition, and elongation with ECM degradation. Given tetherin’s role in regulating MT1-MMP in the pericellular environment, we propose that tetherin affects invadopodia elongation and their ability to degrade the ECM.

It remains to be determined how *Bst-2*/tetherin expression is upregulated in cancer cells. Tetherin expression can be induced by IFNs (Neil et al., 2007) (Fig. S4, A and B) highlighting that IFN and inflammatory phenotypes in the tumor microenvironment may trigger tetherin expression with a predicted impact on regulating the balance of exosome release and retention. Additionally, it is possible that tetherin influences ECM degradation and breast cancer cell migration through mechanisms beyond MT1-MMP and exosome tethering. For example, tetherin can interact with the transcription factor NF-κB (Tokarev et al., 2013) which in turn regulates the expression of several MMPs (Bond et al., 2001) and invadopodia formation (Hu et al., 2022).

## Materials and methods

### Cell culture, transfection, and drug treatments

MDA-MB-231 cells were purchased from Caltag Medsystems (UK). MDA-MB-231 cells were grown in DMEM supplemented with 10% (v/v) FBS, L-glutamine, penicillin-streptomycin, and maintained at 37°C in a 5% (v/v) CO_2_ incubator. For invadopodia formation assay, MDA-MB-231 cells were grown in L-15 supplemented with 15% (v/v) FBS, penicillin-streptomycin, and maintained at 37°C in a 1% (v/v) CO_2_ incubator.

MDA-MB-231 cells were transfected using TransIT®-BrCa Transfection reagent (Mirus) according to the manufacturer’s protocol.

HEK293T cells were purchased from Merck. HEK293T cells were grown in DMEM supplemented with 10% (v/v) FBS, L-glutamine, penicillin-streptomycin, and maintained at 37°C in a 5% (v/v) CO_2_ incubator. HEK293T cells were transfected using TransIT®-293 Transfection reagent (Mirus) for the generation of retrovirus and lentivirus.

Cells were treated with 100 nM Bafilomycin A1 for 16 hours, or with 50 μg/mL Leupeptin for 2h or 4h, or with 1000U/mL Interferon-α (IFN-α) for 16 hours.

### Generation of cell lines

Bst-2 KO cells were generated using a lentiviral CRISPR/Cas9 approach. Cas9 stable MDA-MB-231 cells were generated by transduction with lentiviral pHR-SIN- pSFFV-FLAG-Cas9-pPGK and selected with 200 μg/mL hygromycin B for 7 days. Bst-2 KO cells were generated by transduction of Cas9-stable MDA-MB-231 cells with lentiviral pKLV-u6gRNA-pGK-Puro2A-BFP and subsequent selection with 1 μg/mL puromycin for 4 days. Bst2-HA rescue cells were generated by transducing Bst-2 KO MDA-MB-231 cells with pLXIN-Bst2-HA and selection with 400 μg/mL G418 for 7 days.

Cell clones were obtained by single-cell culture in 96-well plates. Mixed cell populations were obtained by sorting with an Aria III cell sorter (BD), upon staining with a PE-conjugated anti-Tetherin antibody (RS38E, Biolegend).

Stable WT MDA-MB-231 expressing either MT1-MMP-pHluorin or CD63-pHluorin were generated by transduction with lentiviral TetOne-MT1-MMP-pHluorin and TetOne-CD63-pHluorin and selected with 1 μg/mL puromycin for 4 days. Expression of MT1-MMP-pHluorin or CD63-pHluorin was induced by the addition of 20 ng/mL doxycycline for 24 hours.

### Plasmids

CD63-pHluorin-SPORT6, VAMP-2-pHluorin-SPORT6 were a kind gift from Dr. Frederick Verweij and Prof Michiel Pegtel (VUmc, Amsterdam, Netherlands). MT1- MMP-pHluorin-pcDNA3.1 was a kind gift from Prof Philippe Chavrier (Institut Curie, Paris, France). Bst-2-HA cDNA was a kind gift from Prof. George Banting (University of Bristol, UK) and was cloned to pLXIN plasmid by restriction digest.

MT1-MMP-pHluorin-TetOne and CD63-pHluorin-TetOne plasmids were generated by inserting MT1-MMP-pHluorin or CD63-pHluorin into a lentiviral TetOne vector by restriction digestion.

All plasmids were confirmed by Sanger sequencing.

### Antibodies

Antibodies were obtained from the following sources: MT1-MMP (ab51074, rb, abcam) (1:100 for IF, 1:500 for WB, 1:10 for cryo immunoEM, 1:50 for surface TEM), MT1-MMP (LEM-2/15.8, ms, Millipore) (1:100 for IF), EEA-1 (C45B10, rb, Cell Signaling) (1:100 for IF), CD63 (H5C6, ms, BioLegend) (1:200 for IF, 1:500 for WB), LBPA (6C4, ms, Millipore) (1:200 for IF), Rab7 (D95F2, rb, Cell Signaling) (1:50 for IF), LAMP-1 (H4A3, ms, BioLegend) (1:200 for IF), ALIX (3a9, ms, GeneTex) (1:500 for WB), syntenin (ab133267, rb, abcam) (1:500 for WB), GAPDH (14C10, rb, Cell Signaling) (1:1000 for WB), calnexin (N3C2, rb, GeneTex) (1:1000 for WB), CD9 (MM2/57, ms, ThermoFisher Scientific) (1:500 for WB), VPS35 (sc-374372, ms, Santa Cruz Biotechnology) (1:200 for IF), SNX27 (ab77799, ms, abcam) (1:200 for IF), Tetherin (ab243230, rb, abcam) (1:200 for IF, 1:3000 for WB), Cas9 (7A9, ms, BioLegend) (1:2000 for WB), HA antibody (3F10, rat, Roche) (1:200 for IF), collagen ¾ (0217-050, rb, ImmunoGlobe) (1:100 for IF), cortactin (4F11, ms, Merck) (1:200 for IF), PE-conjugated anti-Tetherin (clone RS38E, ms, BioLegend) (1:200 for flow cytometry), HRP-conjugated anti-rabbit or anti-mouse (1:10000 for WB), AlexaFluor 488,555 -conjugated anti-rabbit, anti-mouse or anti-rat (1:200 for IF) from ThermoFisher Scientific, Protein A conjugated to 5 or 10 nm gold particles (PAG; 1:50) were purchased from Cell Microscopy Center, Utrecht University Hospital, Utrecht, Netherlands.

### Reagents

CF-488 or -633 phalloidin was from Biotium and used 1:200 for IF. Gelatin-FITC was from BioVision. Cathepsin B Assay Kit (Magic Red) (ab270772) was purchased from abcam and used according to the manufacturer’s protocol. Rat tail collagen was from Corning (Merck) and was conjugated in house with AlexaFluor-647 (A20006, ThermoFisher Scientific). Leupeptin hemisulfate, Bafilomycin A1 and Doxycycline were purchased from Cayman Chemical Company. Recombinant human Interferon-α was purchased from Sigma-Aldrich.

### sEV isolation

sEVs were prepared from conditioned media incubated for 48h on sub-confluent cells (10 million cells). FBS supplemented medium was previously centrifuged at 100,000 x *g* for 16h to remove FBS-derived EVs. Conditioned media were centrifuged at 300 x *g* (15 min, 4° C) and 2000 x *g* (20 min, 4°C) to remove cell debris, and concentrated by ultrafiltration on a 10,000 MWCO filter (Amicon, Merck-Millipore) to obtain a concentrated conditioned media (CCM). sEVs were then isolated from the CCM by size exclusion chromatography (SEC) (IZON Science). Fractions from 7 to 10 were collected (EV enriched fraction), pooled, and concentrated again using a 10,000 MWCO filter (Amicon, Merck-Millipore). Fractions from 13 to 15 were collected as “soluble protein enriched fraction”. Samples were either used right after isolation or stored at -80°C.

### Western Blot

Cells were lysed on ice in lysis buffer (20 mM Tris, 150 mM NaCl, 1% Triton X-100, and 1 mM EDTA, pH 7.2) supplemented with a protease inhibitor cocktail (Roche). Cell lysates or isolated sEVs were incubated with sample buffer with or without 350 mM 2-mercaptoethanol (Sigma-Aldrich), incubated at 60°C for 30 min, loaded on 4– 12% Bis-Tris gels (Nu-PAGE, Invitrogen), and transferred on nitrocellulose membranes (GE Healthcare). Membranes were blocked in 0.1% Tween/PBS (PBS/T) with 5% non-fat dried milk, incubated with indicated primary (overnight, 4°C) and secondary antibodies (1 hour, room temperature) diluted in PBS/T-milk. Western blots were developed using the SuperSignal West Pico PLUS chemiluminescent substrate (ThermoFisher Scientific). The presented immunoblots are representative of at least three independent experiments. Signal intensities were quantified with Image J Fiji software. Quantifications were from at least three independent experiments.

### NanoFCM

sEVs were analyzed using the Flow Nano Analyzer (NanoFCM), according to the manufacturers protocol (Tian et al., 2020). Briefly, lasers were calibrated using 200 nm control beads (NanoFCM Inc.), which were analyzed as a reference for particle concentration. A mixture of various sized beads (NanoFCM Inc.) was analyzed to set a reference for size distribution. PBS was analyzed as background signal. Particle concentration and size distribution were calculated using NanoFCM software (NanoFCM profession V1.0) and normalized to the dilution necessary for adequate NanoFCM-reading (usually 1:200 to 1:500).

### Flow cytometry

Cells were gently trypsinized and centrifuged at 300 x *g* in a benchtop centrifuge at 4 °C for 5 min. Subsequently, cells were blocked in 1% BSA/ PBS for 30 min on ice.

Surface staining of cells was performed in PBS with 0.5% BSA + 1 mM EDTA (FACS buffer) for 30 min on ice. Cells were then stained with PE-conjugated anti-tetherin antibody in FACS buffer and washed in cold PBS twice. Fluorescence intensity was measured on a four laser Cytoflex S (Beckman coulter, 488 nm, 640 nm, 561 nm, 405 nm).

### Electron microscopy

For conventional electron microscopy, cells were plated to Thermanox (Nunc) coverslips, fixed with 4% PFA /0.1M cacodylate buffer for 30 minutes, and washed with 0.1M cacodylate buffer. Cells were stained using 1% osmium tetroxide/1.5% potassium ferrocyanide for 1 hour, washed, dehydrated with ethanol, and infiltrated with Epoxy propane (CY212 Epoxy resin:propylene oxide) before being embedded in Epoxy resin. Epoxy was polymerized at 65°C for 24h before Thermanox coverslips were removed using a heat-block. 70nm sections were cut using a Diatome diamond knife mounted to an ultramicrotome. Ultrathin sections were stained with uranyl acetate and lead citrate.

To enable luminal surface epitopes to be labelled, cells were fixed with 4% PFA/PBS, washed with 50 mM glycine/ PBS before being blocked with 1% BSA/PBS. Coverslips were inverted over drops of rabbit anti-MT1-MMP for 1 hour and washed before being incubated with protein A gold (PAG) 10 nm for 30 minutes. Following gold labelling, cells were re-fixed using 2% PFA/ 2.5% glutaraldehyde/ 0.1 M cacodylate before being processed for conventional electron microscopy as described above.

For ultrathin cryosections and immunogold labeling, cells were grown on flasks and fixed with 2% PFA/ 0.125% glutaraldehyde/ 0.1 M phosphate buffer. Cell pellets were embedded in 10% gelatin and infused in 2.3M sucrose. Gelatin blocks were frozen and processed for ultracryomicrotomy. Ultrathin sections were immunogold labeled using anti-MT1-MMP and PAG 10 nm as previously described (Slot and Geuze, 1985).

For sEV negative staining, EV samples were spotted on formvar/carbon-coated copper/palladium grids for 20 minutes and fixed with 2% PFA/PBS for 20 minutes. Grids were washed 5 times with water and negative staining was performed using uranyl acetate 0.4% in methylcellulose for 10 minutes.

A FEI Tecnai transmission electron microscope at an operating voltage of 80kV was used to visualize samples, mounted with a Gatan US1000 CCD camera (2K x 2K; 4 MegaPixel).

### Immunofluorescence and microscopy

Cells were grown on coverslips and fixed for 20 min at RT with 4% PFA/PBS and quenched for 10 min with 50 mM glycine/PBS. Cells were incubated with 0.2% BSA/0.1% saponin/PBS before incubation with primary and secondary antibodies in the same buffer.

For cell surface labelling, cells were grown on coverslips, incubated with 1%BSA/PBS before incubation with primary antibodies in the same buffer on ice for 45 min. Cells were fixed for 20 min at RT with 4% PFA/PBS, quenched for 10 min with 50 mM glycine/PBS, and incubated with secondary antibodies.

For colocalization of MT1-MMP or CD63 with Magic Red, MDA-MB-231 WT expressing either MT1-MMP-pHluorin or CD63-pHluorin were incubated with 1X Magic Red for 20 minutes according to the manufacture’s protocol. Cells were fixed with 4% PFA/PBS for 15 minutes, washed with PBS, and imaged immediately.

Incubation with CF-conjugated phalloidin was performed during the secondary antibody incubation. Cells were washed using the same buffer and coverslips were mounted using Prolong Gold antifade reagent with DAPI (Invitrogen). Cells were imaged using a LSM700 confocal microscope (63×/1.4 NA or 40×/1.3NA oil immersion objective; ZEISS) or a LSM780 microscope (63×/1.4 NA or 40×/1.3NA oil immersion objective; ZEISS).

### Gelatin degradation assay

FITC-conjugated gelatin coated coverslips were prepared as previously described (Díaz, 2013). Briefly, sterile coverslips were coated with a pre-heated gelatin mix (5mg/mL unstained gelatin +1 mg/mL FITC-gelatin in 2% sucrose) for 30 minutes at RT. Coated coverslips were fixed with 0.5% glutaraldehyde for 15 minutes on ice, incubated with 5mg/mL Sodium Borohydride for 15 minutes at RT and washed three times with PBS. Coverslips were equilibrated with serum-containing media for at least 1h at 37°C before adding the cells. 50,000 cells were seeded on gelatin coated coverslips and incubated for 5 hours before fixation with 4% PFA/PBS and quenched for 10 min with 50 mM glycine/PBS.

Cells were incubated with 0.2% BSA/0.1% saponin/PBS before incubation with phalloidin-CF633. 0.3 μm Z-stacks were acquired using a LSM700 confocal microscope (63×/1.4 NA or 40x/NA oil immersion objective; ZEISS).

For quantification using ImageJ Fiji, a max projection of 3 Z-stacks was created and a manual threshold was used to segment and measure the area of degraded gelatin (black areas in the fluorescent gelatin) for each cell. Data are shown as relative area of degraded gelatin and values have been normalized to the WT condition. At least 40 cells per condition from 5 biologically independent experiments were analyzed.

### Pericellular collagenolysis assay

2.5 x 10^5^ cells were trypsinized and resuspended in 200 μl of 2.2 mg/mL AF-647 collagen-I mix. 40 μl of cell-collagen suspension was loaded into each well of a 96 well plate to make a 3D gel (at least 4 gels per condition). 3D gels were let to polymerize for 30 minutes at 37°C before adding full media and incubating them for 24 hours. 3D gels were fixed with 4% PFA/PBS for 30 minutes at 37°C before blocking with 0.2% BSA/0.1% saponin/PBS and staining with collagen ¾ rabbit antibody to detect cleaved collagen (1:100 for 2 hours at 4°C) and phalloidin-CF488.

0.3 μm Z-stacks were acquired using a LSM700 confocal microscope (40x/1.3NA oil immersion objective; ZEISS) or a LSM780 microscope (40x/1.3NA oil immersion objective; ZEISS). For quantification using ImageJ Fiji, a max projection of 7 z-stacks was created and a manual threshold was used to segment and measure the areas positive for collagen ¾ staining. The number of cells per picture was counted manually and used to determine the average area of pericellular collagenolysis per cell. In total, at least 300 cells per condition were analyzed.

### Invadopodia formation assay

Coverslips were coated with a thin layer of 2.2 mg/mL AF-647 collagen-I mix and collagen was let to polymerize for 3 minutes at 37°C. 5 x 10^4^ cells, resuspended in 1mL of media were seeded. Cells were incubated for 1 hours at 37°C to allow invadopodia formation. Cells were incubated with 4% PFA/ 0.5% Triton-X-100/PBS for 90 seconds, fixed with 4% PFA/PBS for 20 minutes, quenched for 10 min with 50 mM glycine/PBS, blocked with 0.2% BSA/0.1% Triton-X-100/PBS and stained with phalloidin-CF488 and anti-cortactin antibody.

0.3 μm z-stacks were acquired using a LSM700 confocal microscope (63×/1.4 NA oil immersion objective; ZEISS).

For quantification using ImageJ Fiji, a max projection of 4 z-stacks was created and a manual threshold was used to segment and measure the area of each cell covered by invadopodia (cortactin positive structures forming along collagen fibrils). At least 50 cells per condition from at least 3 biologically independent experiments were analyzed.

### TIRF microscopy

TIRF microscopy of cells expressing pHluorin tagged proteins was performed as previously described (Bebelman et al., 2020). Cells were grown on glass bottom 35 mm dishes and transfected with either CD63-pHluorin-SPORT6 or MT1-MMP- pHluorin-pcDNA3.1 or VAMP-2-pHluorin-SPORT6 as described above. An Elyra 7 (ZEISS) with an alpha Plan Apochromat 63×/1.46 Oil Korr M27 TIRF objective was used. Images were acquired at 2 or 3 Hz for 2 minutes. Live-imaging experiments were performed in culture medium at 37°C and 5% CO2.

Movies were analyzed manually for the presence of MVE- plasma membrane fusion events, defined as a sudden increase in fluorescence intensity. Fusion activity was defined as the number of fusion events per μm^2^ of cell surface. The duration of a fusion event was defined as the time intercurrent between the peak intensity and the moment when the fluorescence intensity has reached back the surrounding background fluorescence.

At least 7 cells from at least 3 biologically independent experiments were analyzed.

### Image analysis

Image analysis methods not described above were performed as follows using ImageJ Fiji version 2.3.0.

#### Pearson’s correlation coefficient

Pearson’s correlation coefficient between two channels was quantified using JACoP plugin of ImageJ Fiji software. At least 25 cells from 3 biologically independent experiments were analyzed.

#### Line scan analysis for MT1-MMP, VPS35 and SNX27 endosomal localization

A line of 3 μm was drawn on endosomes positive for MT1-MMP and VPS35 (or SNX27) using ImageJ Fiji. Fluorescence intensity pixel grey values were measured using the Plot Profile function and normalized as percentage of the maximum value measured along the line.

At least 50 endosomes from at least 12 cells were analyzed.

#### Overlap ratio of MT1-MMP or CD63 with Magic Red

A circle was drawn around each MT1-MMP or CD63 positive compartment using ImageJ Fiji and saved as an ROI. Each ROI was scored for the presence or absence of Magic Red signal. The number of Magic Red positive ROIs was counted and its ratio against the total number of ROIs was calculated and defined as overlap ratio with Magic Red. At least 20 cells from 2 biologically independent experiments were analyzed.

#### Number of tethered sEVs at the cell surface

sEVs in proximity of the plasma membrane (within 500 nm from the membrane) were counted manually. The length of the plasma membrane (μm) was measured using ImageJ Fiji software. Single cells were analyzed and the ratio number of sEV/ length of the plasma membrane was calculated. At least 35 cells from 3 biologically independent experiments were analyzed per condition.

### Statistical analysis

The number of analyzed experiments and cells is indicated in the figure legends or above in the methods. No samples were excluded from the analysis. All statistical analyses were performed using GraphPad version 10.

We primarily tested data for normal distribution using Anderson-Darling, D’Agostino and Pearson, and Shapiro-Wiki, and Kolmogorov-Smirnov tests. For parametric data, either unpaired or paired *t*-test (for comparison between two datasets) or Ordinary one-way ANOVA (for comparison of more than two datasets) were performed. For non-parametric data, Kruskal-Wallis test was performed. All data are sown as mean ±SEM. Significant differences are indicated as follows: ****, P <0.0001; ***, P < 0.001; **, P < 0.01; *, P < 0.05; n.s., P ≥ 0.05. Only P < 0.05 was considered as statistically significant.

## Data availability

All data generated or analyzed during this study are available in the article and its online supplemental material. Requests for materials should be addressed to J. R. Edgar.

## Acknowledgments

We wish to thank the microscopy and flow cytometry facilities at the Department of Pathology, University of Cambridge. We also wish to thank the electron microscopy facility at Cambridge Institute for Medical Research, University of Cambridge, and the Cambridge Advanced Imaging Centre, University of Cambridge.

Funding: JRE and RP were supported by a Sir Henry Dale Fellowship jointly funded by the Wellcome Trust and the Royal Society (216370/Z/19/Z). JRE was also supported by a Royal Society Research Grant (RGS/R2/192145).

## Author contribution

R Palmulli: Formal analysis, Investigation, Methodology, Validation, Visualization, Writing – original draft, Writing - review and editing; H.K. Jackson: Formal analysis, Investigation, Writing - review & editing; J. R. Edgar: Conceptualization, Funding acquisition, Methodology, Project administration, Supervision, Validation, Writing - original draft, Writing - review & editing.

---

Figure S1. Supplemental data related to Fig. 1.

**(A)** Confocal analysis of endogenous MT1-MMP localization with VPS35 or SNX27 in MDA-MB-231 WT cells. Scale bars = 10 μm. Dashed lines represent the 3 μm drawn to perform the line scan analysis in (B). **(B)** Line scan analysis of MT1-MMP with VPS35 or SNX27 fluorescence intensity in endosomes. n ≥ 50 endosomes, error bars represent SEM. **(C)** Western blotting analysis of WCL from MDA-MB-231 cells control or treated with Leupeptin for 2 or 4 hours. **(D)** Enrichment of MT1-MMP in WCL from MDA-MB-231 cells control or treated with Leupeptin for 2 or 4 hours. N = 3 biologically independent experiments, paired *t* test, * p= 0.0204. **(E)** Confocal analysis of endogenous MT1-MMP localization with LAMP-1 in MDA-MB-231 cells control or treated with Leupeptin for 4 hours. Scale bars = 10 μm. **(F)** Confocal analysis of endogenous MT1-MMP localization with LAMP-1 in MDA-MB-231 cells control or treated with BafA1 for 16 hours. Scale bars = 10 μm. **(G)** Confocal analysis of MT1-MMP or CD63 localization with active Cathepsin B dye Magic Red in MDA-MB-231 WT cells. Scale bars = 10 μm. **(H)** Overlap ratio of MT1-MMP or CD63 with Magic Red (mean ± SEM). n = 22 for CD63, 25 for MT1-MMP, cells from 2 biologically independent experiments, unpaired *t* test. **(I)** EM micrograph of sEVs isolated from MDA-MB-231 WT cells. Scale bars = 200 nm. **(J)** NanoFCM analysis of sEVs isolated from MDA-MB-231 WT cells. Size distribution of sEVs (mean + SEM), 3 biologically independent experiments. **(K)** EM micrograph of MDA-MB-231 WT cells treated with DMSO or BafA1 for 16 hours. Scale bars = 1 μm.

Figure S2. Supplemental data related to Fig. 3.

**(A)** Confocal analysis of tetherin localization with CD63 or LAMP-1 in MDA-MB-231 cells. Scale bars = 10 μm. **(B)** Pearson’s correlation coefficient of tetherin and CD63 or LAMP-1. n = 44 for CD63, 29 for LAMP-1, cells from 2 biologically independent experiments, error bars represent SEM. **(C)** Confocal analysis of tetherin localization with LAMP-1 in MDA-MB-231 cells control or treated with BafA1 for 16 hours. Scale bars = 10 μm. **(D)** Surface staining of MDA-MB-231 WT cells labelled with tetherin. Scale bars = 10 μm.

Figure S3. Supplemental data related to Fig 4.

**(A)** Confocal analysis of endogenous MT1-MMP localization with different endosome/lysosome markers in MDA-MB-231 cell lines. Scale bars = 10 μm. **(B)** Pearson’s correlation coefficient of MT1-MMP and endosome/lysosome markers. n ≥ 25 cells from 3 biologically independent experiments, mean ± SEM, ordinary one-way ANOVA, * p= 0.0111 and ** p=0.0097 for RAB7, ** p=0.0025 and ** p= 0.0022 for LAMP-1. **(C)** Confocal analysis of endogenous MT1-MMP localization with VPS35 in MDA-MB-231 cell lines. Scale bars = 10 μm. **(D)** Relative enrichment of the protein content in WCL from MDA-MB-231 cell lines (mean ± SEM). n = 5 biologically independent experiments, two-way ANOVA. **(E)** Fusion activity of MT1- MMP -pHluorin. n = 7 cell fusion events per reporter from 3 biologically independent experiments, *t*-test with Welch correction. Signal duration of fusion events of MT1- MMP -pHluorin. n = for WT, 54 for Bst-2 KO, 44 fusion events per reporter from 3 biologically independent experiments, ordinary one-way ANOVA, * p=0.0142.

Figure S4. Supplemental data related to Fig 5.

**(A)** Confocal analysis of MDA-MB-231 WT control or treated with IFN-α or MDA-MB- 231 WT + Bst-2-HA stained for tetherin. Scale bars = 10 μm. **(B)** Western blotting analysis of tetherin expression in MDA-MB-231 cell lines control or treated with IFN-α. **(C)** Confocal analysis of collagen degradation assay. MDA-MB-231 WT control or treated with IFN-α were embedded in 3D fluorescent collagen gels, incubated for 24 hours, fixed and stained with phalloidin and for the cleaved collagen new epitope using anti-Col-3/4 antibody. Scale bars = 10 μm. **(D)** Quantification of pericellular collagen degradation expressed as average degraded collagen area per cell by MDA-MB-231 WT control or treated with IFN-α (mean ± SEM). n = 36 for WT, 57 for WT+ IFN-α confocal images (≥ 300 cells) from 3 biologically independent experiments, Kruskal-Wallis test.

Video 1. TIRF-M fusion event examples of CD63-pHluorin (left), VAMP-2- pHluorin (middle), and MT1-MMP-pHluorin (right) in WT MDA-MD-231 cells.

Video 2. TIRF-M fusion event examples of CD63-pHluorin in WT (left), +Cas9 (middle) and Bst-2 KO (right) MDA-MB-231 cell lines.

Video 3. TIRF-M fusion event examples of MT1-MMP-pHluorin in) MDA-MB-231 cell lines.

