## Supplementary figures and images for "Tethered exosomes containing MT1-MMP contribute to extracellular matrix degradation during breast cancer progression"

### Supplemental Figures 1-4

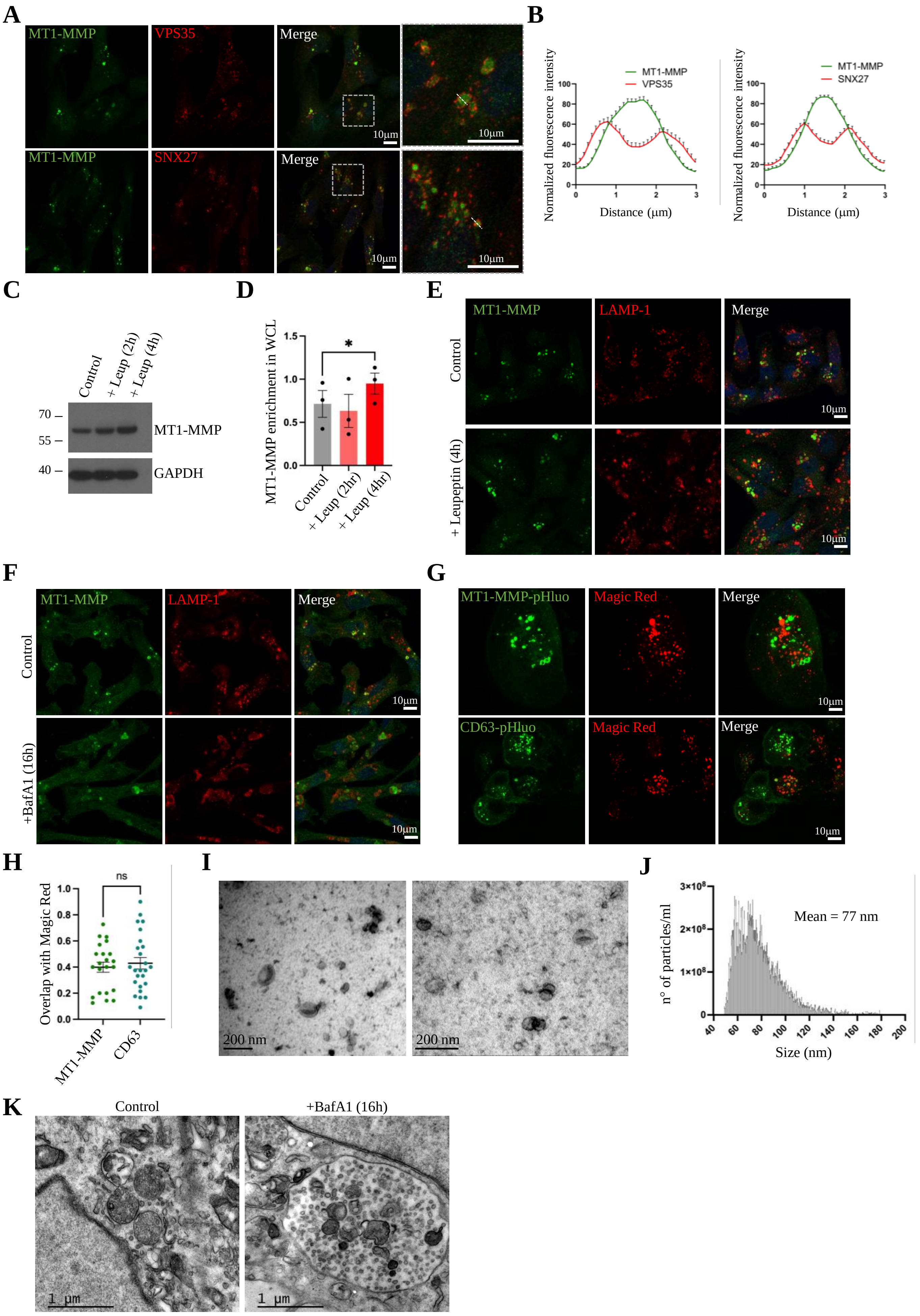

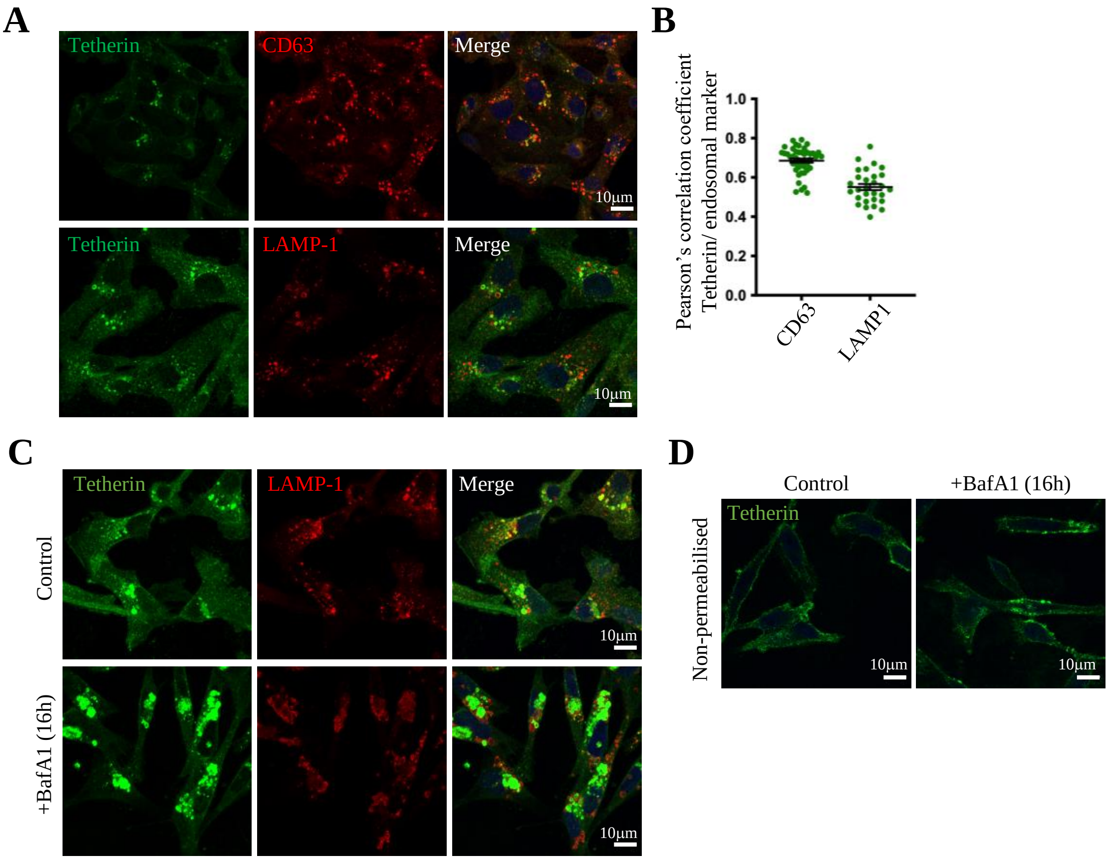

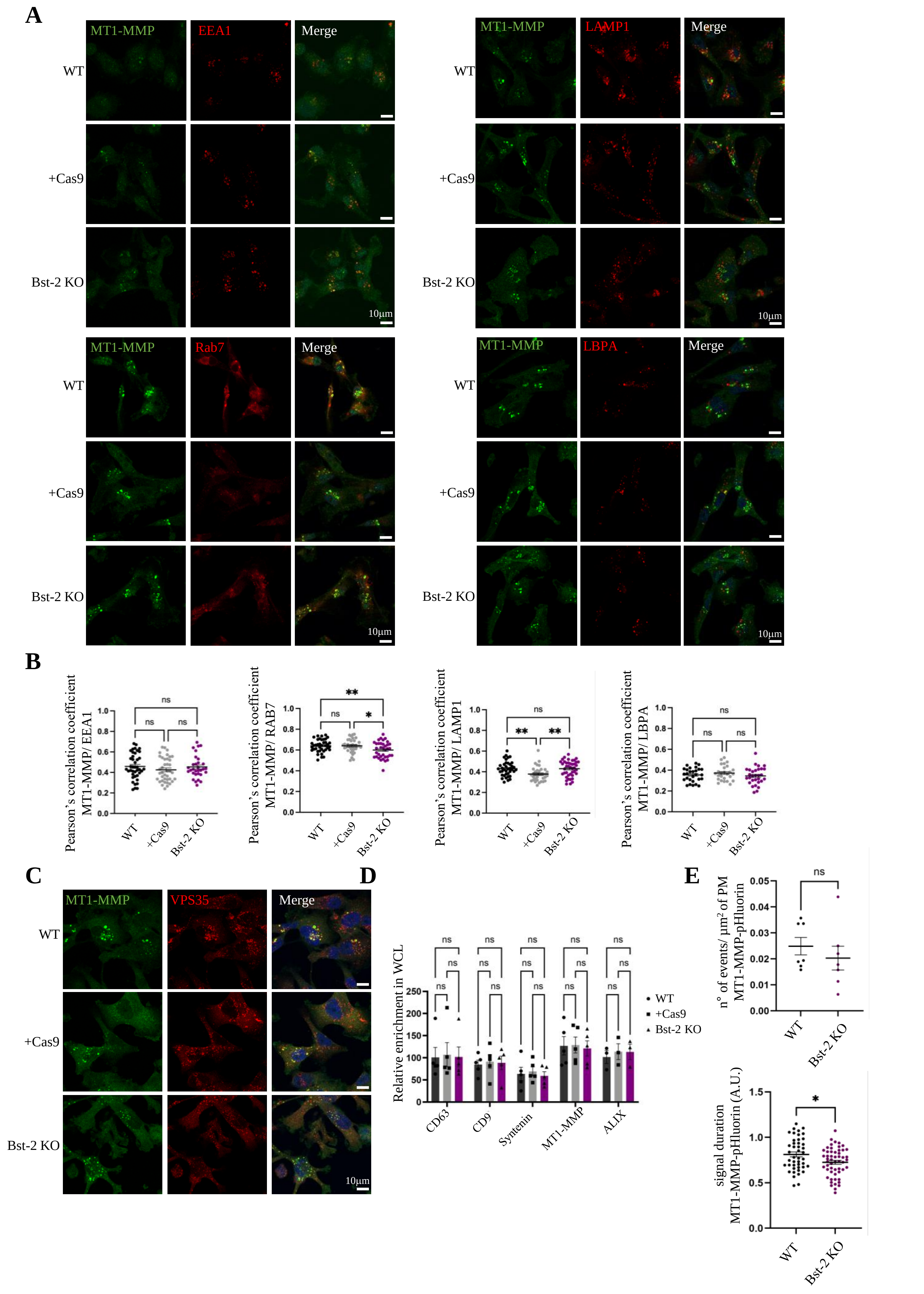

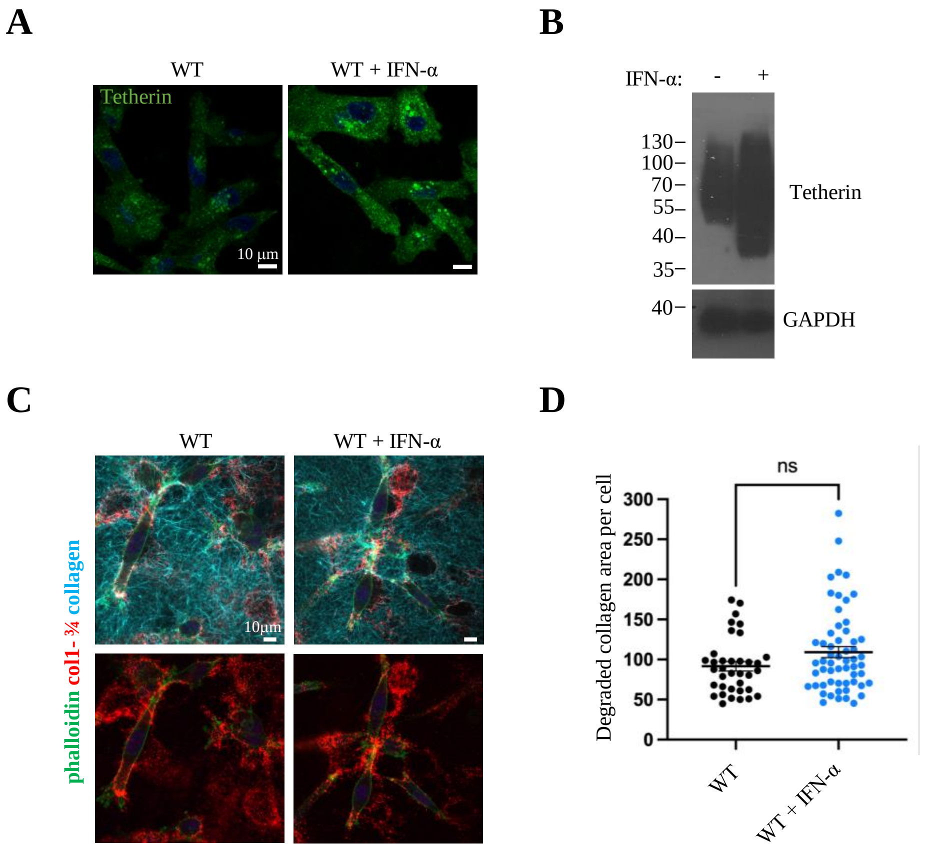
